# The ganglioside GD3 and its Synthase (ST8SIA1) as novel senescence markers associated with osteoarthritis

**DOI:** 10.1101/2025.03.22.640206

**Authors:** Christina Fissoun, Georges Maroun, Raissa Silva, Margot Milano, Benoit Guibert, Louis Dagneaux, Rosanna Ferreira-Lopez, Florence Apparailly, Thérèse Commes, Eric Gilson, Christian Jorgensen, Julien Cherfils-Vicini, Yves-Marie Pers, Jean-Marc Brondello

## Abstract

Osteoarthritis (OA), is the most common age-induced degenerative joint disease. It is associated with synovial inflammation, subchondral bone remodeling and cartilage degradation. One of the significant emerging causes of OA progression is senescent cell accumulation within the joint compartment during lifespan. Currently, there are no therapeutic approaches nor stratification tools that rely on the senescence burden in OA. In this study, we identified the b-series ganglioside 3 (GD3) as new senescent cell surface marker associated with OA. Joint RNA sequencing analysis revealed an increase expression of the GD3 synthase, *ST8SIA1* in cartilage, synovial tissue, and subchondral bone marrow from OA patients compared to healthy donors. Moreover, we revealed a strong correlative association between the expression of *ST8SIA1* and GD3 production with senescence hallmarks in an *in vitro*-induced 3D organotypic OA cartilage model but also with cartilage histological grading scores in human and preclinical murine OA joints. Anti-GD3 cell sorting showed that GD3-positive human OA chondrocytes or human OA synoviocytes are enriched in senescence and SASP markers compared to GD3-negative counterparts confirming that GD3 is a cell surface marker linked to the senescence stage. Intra-articular anti-GD3 antibody delivery in experimental OA model, reduced local expression of senescence and OA markers in association with a protection against OA-induced subchondral bone remodeling. Our research demonstrates a compelling linkage between *ST8SIA1* gene, GD3 and senescence in OA pathology, revealing knowledge and perspectives for a better understanding and anti-senescence treatment of OA pathogenesis.

## INTRODUCTION

Osteoarthritis (OA) is the most common chronic degenerative joint disease in elderly population particularly affecting the knees [1]. OA is characterized by progressive cartilage deterioration toward a hypertrophic stage, osteophyte formation, subchondral bone remodeling and low-grade synovial inflammation. Its economic burden is high due to the absence of curative treatments and the aging of the population. In the last decade, researchers have begun to explore a new strategy for treating OA by targeting joint cells that display a phenotype associated with cellular senescence [2]. Cellular senescence is a cell response to tissue injuries and chronic cues [3]. In OA, senescent cells are characterized by high expression of cell cycle inhibitors (e.g. p16^INK4a^, p15^INK4b^ or p21) and the presence of the senescence-associated secretory phenotype (SASP) that includes catabolic (e.g. MMP13, ADAMTS4/5), inflammatory (IL-11, IL17D, S100A4) and trophic factors (GREM1, TGFb1) [3,4]. The pharmacogenetic clearance of these senescent cells has beneficial effects in preclinical murine models of OA by reducing joint impairment and disease progression [5]. However, to date, no senotherapeutic strategy has shown positive results in clinical trials for the treatment of OA patients, possibly due to the insufficient knowledge and characterization of the role of cell senescence in the disruption of joint homeostasis and OA progression.

Gangliosides, a family of glycosphingolipids, play a critical role in normal neuronal development, complement system regulation but also nerve and bone tissue maintenance [6,7]. GD3, a ganglioside produced by the synthase ST8SIA1 from GM3 precursor, has thus emerged as an important player in various cellular functions and disease mechanisms [7] including cancer [8]. Interestingly, GD3 KO mice harbor an increase in bone mass with aging compared to wild-type mice [9]. In addition, the GD3 precursor GM3 has been implicated in chondrocyte hypertrophy during cartilage remodeling following focal injury [10]. Genetic inhibition of GM3 synthesis has indeed been shown to improve cartilage repair, suggesting a potential role for GM3 and GD3 in osteo-articular homeostasis [9] [10]. Finally, we recently discovered that GD3 and its synthase is linked to replicative senescence [11] and represent a novel senescence immune checkpoint (SIC) that regulates NK cell-mediated senescent cell clearance [11]. In aged mice, we observed that anti GD3 immunotherapy administrated intra-peritoneally was sufficient to improve aged associated bone remodeling [11]This discovery highlights a potential broader role for GD3 in several senescence-driven pathologies.

In this study, we used human OA joint tissue biobanks, an *in vitro*-induced 3D organotypic model of senescent cartilage and *in vivo* immuno-targeting approaches in preclinical murine OA model to investigate the association of GD3 and its synthase with OA-related senescence.

## MATERIALS AND METHODS

### Human Cartilage Samples

After signature of the informed consent, human OA cartilage and synovial sample for banking were obtained from patients with OA undergoing knee joint replacement surgery at the Lapeyronie hospital, Montpellier, France. Sample collection and banking were approved by the national and local ethical committees (IRB-MTP_2022_11_202201238).

### Cell Culture

Knee cartilage samples were incubated in 2.5mg/mL of pronase (# 1074330001, Sigma Aldrich) at 37°C for 1 hour and then in 2 mg/mL of collagenase II (# C6885, Sigma Aldrich) at 37°C overnight. Synovial samples were incubated in 1mg/mL of collagenase IV (# C4-28, Sigma Aldrich) at 37°C overnight. Digested samples were filtered through a 70 μm strainer, then the isolated cells were cultured in D-MEM/1% penicillin-streptomycin/1% L-glutamine/10% fetal bovine serum, at the density of 30,000 cell/cm^2^ until confluency. Osteoclasts were differentiated *in vitro*. Monocytes/ macrophages were isolated from the bone marrow of 3-month-old WT male mice, either NaCl either CiOA group, by flushing cells from the bone marrow of femora and tibiae. The flushed bone marrow cells were cultured in 6-well plates, with osteoclastogenesis medium, containing αMEM Glutamax, with 10% fetal bovine serum, 50 μM 2-mercaptoethanol, 1% penicillin-streptomycin solution, 25 ng/mL M-CSF (Miltenyi 130-101-705) and 50 ng/mL RANKL (Peprotech 315-11C) for 4 days and 10 days.

### Immunohistochemistry

Joint samples were embedded in paraffin and cut into sections (3 µm). Sections were deparaffinized in xylene and rehydrated in ethanol. Then, tissue sections were incubated with pepsin (DAKO, 40 mg/mL) for 30 min for antigen retrieval, permeabilized with 0.3% Triton X-100 in PBS/3% BSA for 3 min, 1% H_2_0_2_ for 10 min followed by incubation in PBS/3% BSA for 30 min. Sections were then incubated with primary antibodies against GD3 (1:200, #MAB2053 Merck), p15^INKb^ (1:200, #orb213719 BIORBYT), p16^INK4a^ (1:200, #250804 ABBIOTECH), p21 (#2947, Cell Signaling) and MMP13 (#ab219620, Abcam) at 4°C overnight. Sections were rinsed with PBS, followed by the Polink-2 Plus HRP Broad Spectrum DAB Detection Kit (#D41-18, Diagomics). Images were taken with a Nanozoomer Digital Pathology System (Hamamatsu) and the number of positive cells was quantified with the software QuPath.

### Magnetic-Activated Cell Sorting (MACS)

Isolated cells (chondrocytes or synoviocytes) were washed twice with PBS and resuspended in PBS, 2 mM EDTA, 0.5% BSA. Cell were incubated on ice with 1μg/μL anti-GD3 monoclonal antibody for 30 min, followed by anti-mouse IgG magnetic beads MACS (#130-048-402 Miltenyi Biotec) on ice for 15 min. Magnetic separation was performed using an LS-positive selection column. The GD3-enriched and GD3-depleted populations were then harvested and subsequently used for RNA extraction.

### Flow Cytometry

Freshly isolated chondrocytes were cultured for 3 days to remove particles from the extracellular matrix and avoid autofluorescence. After trypsinization, chondrocytes were incubated on ice with 1μg/μL anti-GD3 monoclonal antibody 30 for min, followed by incubation on ice with a secondary anti-mouse antibody (APC-Alexa Fluor) for 30 min. After washing, DAPI was added to exclude dead cells. Acquisition was performed with a BD FACSCanto II flow cytometry system (BD Biosciences) and analysis with the Kaluza software. Following synovial digestion as described, freshly isolated synoviocytes were immediately incubated at room temperature with a viability marker to exclude dead cells. After washing with phosphate-buffered saline (PBS), the cells were incubated on ice with 1 µg/µL anti-GD3 monoclonal antibody for 30 minutes, followed by a further 30-minute incubation on ice with a secondary anti-mouse antibody (APC-Alexa Fluor). Acquisition was performed with the Cytek Aurora spectral cytometry system (Cytek Biosciences) and analysis with the FlowJo software.

### Gene Expression Analysis by RT-qPCR

For gene expression experiments, total RNA was extracted from ground frozen joint samples using the Phenol/Chloroform method. RNA quality was checked by spectral analysis (A260/ 280 nm), and then samples were stored at −80°C. Reverse transcription (RT) was performed using the M-MLV reverse transcriptase (Invitrogen; 28025013; 5U/μL final concentration), 500ng total RNA, a random hexamer primer (Thermo Scientific, GER; S0142; 10ng/μL final concentration), and dNTPs (Roche, CH; 1 277 049; 5mM final concentration) in M-MLV reverse transcriptase buffer (Invitrogen; 18057-018), for a total volume of 20μL. SYBR Green-based quantitative PCR (qPCR) was performed using the LightCycle® 480 SYBR Green I master reaction mix (Roche, 04707516001), 10 ng of cDNA, and the LightCycler 480 real-time PCR system (Roche) (40 cycles of amplification). Raw data (Ct values) were analyzed using the comparative Ct method. Gene expression data were calculated as relative to the expression of the housekeeping gene *Rpl0* (2^−deltaCT^ method).

### Data Collection and Bioinformatics

For cartilage bulk RNA-seq data, public RNA-seq datasets were recovered from https://www.ebi.ac.uk/arrayexpress/experiments/E-MTAB-6266. The datasets included samples from 60 patients with OA following total knee replacement and 10 control non-OA patients following amputation. A new methodology based on a k-mer approach was used to quantify the gene expression profiles. The automated selection of specific k-mers was ensured by the Kmerator tool (https://github.com/ Transipedia/kmerator). Then, k-mers were directly quantified in the indexed fastq files using the Transpedia website (https://Transipedia.fr). The quantification was expressed as the average count of all k-mers for one transcript, normalized by millions of total k-mers in the raw file. The heatmap was created on R with the ggplot2 package. For synovial single-cell RNA-seq data, public RNA-seq data were recovered from the source paper [12]. Fastq files were downloaded from NCBI GEO GSE15280. In total, 10,640 synovial tissue cells were counted. Cells were grouped into an optimal number of clusters using Seurat. For cluster annotation, the CellDex R package was used. The top upregulated genes for the three synovial cell sub-clusters were defined using data from the following paper [13].

### Animal Experiments

12-week-old C57BL/6JCJ male mice were obtained from Janvier Laboratory. Animal experiments were performed in accordance with the guidelines by the local ethics committee on animal research and care (1872-32846). After sacrifice, the knee joints were isolated, fixed in 4% paraformaldehyde at 4°C for 2 days (immunohistochemistry analysis) or for 7 days (bone and cartilage morphometric analysis).

### Experimental OA and Intra-Articular Injection of Antibodies

Experimental OA was induced in 12-week-old male mice according to the CiOA procedure. Under general anesthesia (isoflurane inhalation), the knee area was disinfected with ethanol and a small cut (few millimeters) was performed in the cutaneous and subcutaneous tissue to visualize and access the knee. Then, each mouse received an intra-articular injection of 5μL (1IU) collagenase VII in the left knee to induce OA and 0.9% NaCl in the right knee as control, followed by a second injection two days later. Intra-articular injections of anti-GD3 (clone R24) or isotype (anti-IgG3) antibodies were performed in mice following the same procedure: 30 μg in PBS of anti-GD3 or isotype antibodies in the left knee at day 7, followed by a second injection 3 days later.

### Safranin-O/Fast Green Staining

For cartilage analysis, mouse knees were fixed in 4% paraformaldehyde at 4°C for 48h, washed in PBS, and then processed for routine histology. Knees were decalcified in 14% EDTA/PBS for 3 weeks and then paraffin-embedded. Tissue sections (5 µm) were rehydrated through a gradient of ethanol and xylene. Sections were then stained with Safranin-O/Fast Green to evaluate cartilage degradation. Cartilage damage was analyzed using an arbitrary score from 0 to 30, based on the OARSI cartilage OA grading system and modified by van den Berg for the assessment of murine knee joints [14] (grading scale of 0–6 for the severity of cartilage destruction and of 0–5 for the extent of damaged cartilage surface).

### Bone Parameter Analyses

Hind leg knees were dissected to remove smooth tissues and scanned in a micro-CT scanner SkyScan 1176 (Bruker, Belgium, 0.5 mm aluminum filter, 45 kV, 500 µA, 18 µm resolution, 0.5° rotation angle). Images were reconstructed using CTAn v1.9, Nrecon v1.6 (Bruker, Belgium) and a three-dimensional (3D) model visualization software program (CTVol v2.0). Misalignment compensation, ring artifacts and beam-hardening were adjusted to obtain the correct re-construction of each paw. Bone degradation was quantified in subchondral bone and in the epiphysis region of the medial and/or lateral plateau for each tibia (CTAn software, Bruker, Belgium). Reconstructed 3D images of joints were obtained using the Avizo software (Avizo Lite 9.3.0, FEI, France).

### Cartilage Structure Quantification by Confocal Laser Scanning Microscopy

Joint cartilage of the tibia medial plateau was scanned through the depth in XYZ-mode with a confocal laser scanning microscope (CLSM; TCS SP5-II, Leica Microsystems, Nanterre, France), a voxel size of 6 μm, a 5× dry objective and a UV laser light source (l¼ 405 nm). Stacks of images were analyzed to quantitatively evaluate several joint cartilage parameters. Cartilage morphometric parameters were assessed in the medial plateau of each tibia using the Avizo software (FEI Visualization Sciences Group, Lyon).

### Statistical analysis

All data are presented as the median or mean ± SEM. The Student’s *t*-test was used for comparisons between experimental groups. For Spearman correlation, the coefficient (*r*) was estimated to determine the linear association between markers. Results were interpreted according to the degree of association as strong (*r* = 0.7–1), moderate (*r* = 0.5–0.7), or low (*r*= 0.3–0.5) after taking significant correlation values into consideration; p-values <0.05 were considered significant. Data were analyzed using the Prism software v10 (GraphPad Software Inc.).

## RESULTS

### *ST8SIA1* mRNA expression increases in OA cartilage and in 3D organotypic OA cartilage model

Based on k-mers method and publicly available RNA-seq data from 10 human healthy cartilage samples (non-OA) and 60 cartilage samples from OA patients obtained following total knee replacement [15] we identified *ST8SIA1* gene, which encodes GD3 synthase, as part of the OA transcriptional signatures **(*Fig. 1a***). *ST8SIA1* was indeed upregulated in OA samples compared to healthy donors as well as *MMP13, ADAMTS4, COL10, IL11* and *IL17D* that are already known OA markers [17–19] **(*Fig. 1b***). Conversely, two mitochondrial sirtuins, *SIRT3* and *SIRT7,* were downregulated in OA samples compared with no-OA samples [20,21]. Interestingly, we did not detect any sex-related difference in the identified signatures although the OA prevalence is higher in women population. To confirm a link between *ST8SIA1* mRNA induced expression and OA pathogenesis, we used our previously published in *vitro* organotypic 3D OA cartilage model. Primary human chondrocytes were placed in micromass culture conditions to form cartilage pellet [23] (***Fig. 1c***). Upon addition of the inflammatory cytokine IL-1β for one week, this *in vitro* 3D model recapitulates OA cartilage phenotypes including senescence-associated SASP production [24]. Indeed, the expression of *CDKN2B, CDKN2A* and *CDKN1A*, three senescence hallmark genes altogether with inflammatory SASP factors (*GREM1, IL1β, IL6*, and *S100A4)* and catabolic SASP factors *(MMP13, ADAMTS5) but also ST8SIA1* were all increased in response to IL-1βtreatment (***Fig. 1d***). Conversely, the expression of the extracellular chondrogenic matrix gene *COL2,* which hallmarks healthy cartilage, was downregulated following IL-1β addition. Finally, on paraffin-embedded slides of this 3D OA model, we could confirm an increase expression of the ST8SIA1 product, the ganglioside GD3 by immuno-histochemistry (IHC) staining using specific antibody in parallel with a decrease in proteoglycan composition revealed by Safranin-O-Fast Green staining. Altogether these findings are in full accordance to RTqPCR results (***Fig. 1e***). Importantly, OA disease is not exclusively associated with cartilage breakdown but also with synovial tissue hypertrophy. We therefore determine whether *ST8SIA1* gene expression is also altered in synovium from OA patients. We thus analyzed publicly available single-cell RNA-seq data from OA synovial membrane samples [12] (***Fig. S1a***). As previously published by others, among the various cell populations identified, we found three different subtypes of fibroblast-like synoviocytes, as well as macrophages/monocytes, endothelial cells, stem cells, muscle cells and other cells belonging to the immune lineage (***Fig. S1b***). The single-cell RNA-seq data showed that *ST8SIA1* was highly expressed in synoviocytes (Uniform Manifold Approximation and Projection, UMAP, plots in ***Fig. S1c***) and specifically in synovial fibroblasts (SF) subtype 2 and subtype 3 (***Fig. S1d-e***). In conclusion, the expression of the GD3 synthase gene, *ST8SIA1*, is enriched in two OA joint compartments.

**Figure 1:**
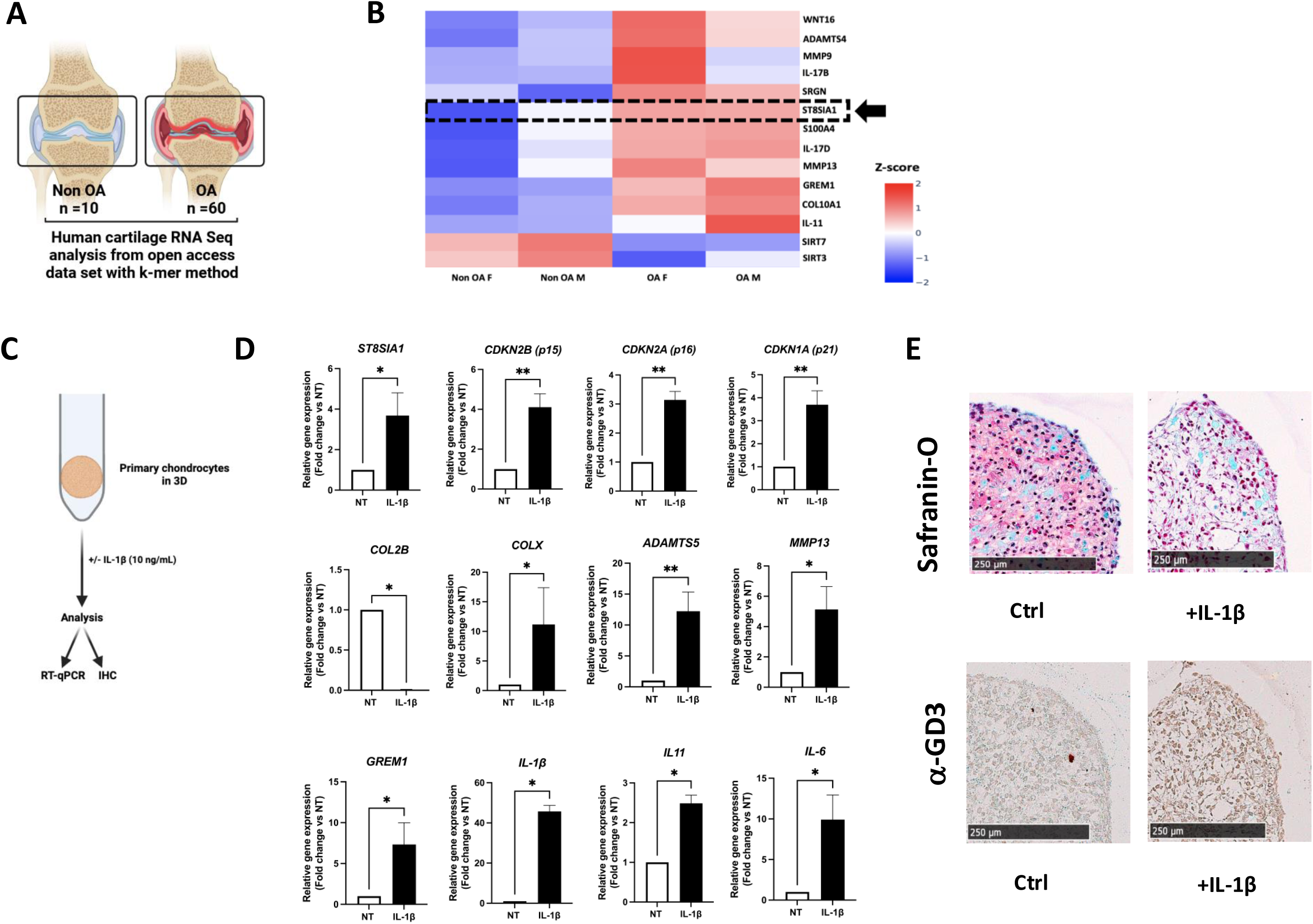
*ST8SIA1 gene* is increased in human OA cartilage and *ST8SIA1 expression* is up-regulated in senescent 3D chondrocytes model. A) Experimental design from RNA-Seq analysis with the help of k-mer method. B) Heatmap represented transcriptomic signature with hallmarks of pathology in cartilage. C) Experiment graphical representation. D) Gene expression analysis of ST8SIA1, senescence, cartilage and hypertrophic markers by RT-qPCR in 3D model chondrocytes treated or not with IL-1β (10ng/ml). mRNA expression levels were normalized to the housekeeping gene RPL0. (n=4) E). Safranin-O staining and GD3 IHC on 3D model chondrocytes treated or not with IL-1β. The scale bar indicates 250 µm. (n=2). *p < 0,05. Statistical significance was determined by paired, two-tailed Student’s t-test.

### Senescent GD3-positive cells are found in OA joint tissue and correlate with cartilage histological OA degrading score

We next asked whether expression level of the ganglioside GD3 was found in OA patient cartilage, correlated with senescence hallmark protein levels altogether with disease progression. To do this, after Safranin-O-Fast Green staining on paraffin slides, we classified collected OA cartilage samples from 10 patients in two groups (low and high score) according to their OA clinical severity based on the Modified Mankin score [25] (***Fig. 2a***). Next, through specific immunohistochemical staining, we quantified the percentages of cells positives for GD3 and the senescence hallmarks p16^INK4a^ or p15^INK4b^ revealing a significant higher level in samples with more severe degradation. The difference was not significant for the percentage of MMP13-positive cells (***Fig. 2b***). Correlation analyses revealed positive associations between the percentage of GD3-positive chondrocytes and the Modified Mankin degrading cartilage score (r=0.7903 p=0.009) as well as with the percentage of p16^INK4a^ (r=0,7091; p=0,0268) and p15^INK4b^ (r=0.8545; p=0,0029) positive cells. Only a non-significant correlation was found with MMP13 positive cells (r=0.6079; p=0.0676) (***Fig. 2c***). We next analyzed by cell sorting the phenotype of GD3-positive chondrocytes in the cartilage from OA patients. For this, we obtained fresh chondrocytes after OA cartilage digestion and stained them with an anti-GD3 antibody (***Fig. 2d*)**. Flow cytometry confirms the presence of GD3 on the OA chondrocytes surface (Supplementary ***Fig. S2a***). Using magnetic-activated cell sorting (MACS) method, we then separated GD3-positive and -negative cells in order to analyze the expression of senescence and OA associated markers in both populations **(*Fig. 2e*).** As expected, *ST8SIA1* expression was higher in GD3-positive than in GD3-negative chondrocytes as was the expression of the senescence markers *CDKN2B* and *CDKN2A* which encode two cell cycle inhibitors p15^INK4b^ and p16^INK4a^ respectively (***Fig. 2f***). Inflammatory SASP factors (*IL6*, *IL8*, *IL1B, GREM1, S100A4* and *IL17D*) as well as SASP catabolic degrading enzymes (*MMP13* and *ADAMTS5*) that drive OA cartilage extracellular matrix breakdown, were also upregulated in GD3-positive chondrocytes. We then looked at OA-associated markers of chondrocyte hypertrophy and found that *COL10*, *RUNX2* and *RUNX3* were also increased in GD3-positive chondrocytes. Conversely, *COL2*, the gene encoding for a functional collagen in healthy chondrocytes, showed lower expression in GD3-positive chondrocytes compared to GD3-negative chondrocytes [22]. Taken together, these data illustrate a connection between the expression of GD3 at the surface of OA chondrocytes and the expression of genes related to senescence and hypertrophy that characterized this joint pathology.

**Figure 2:**
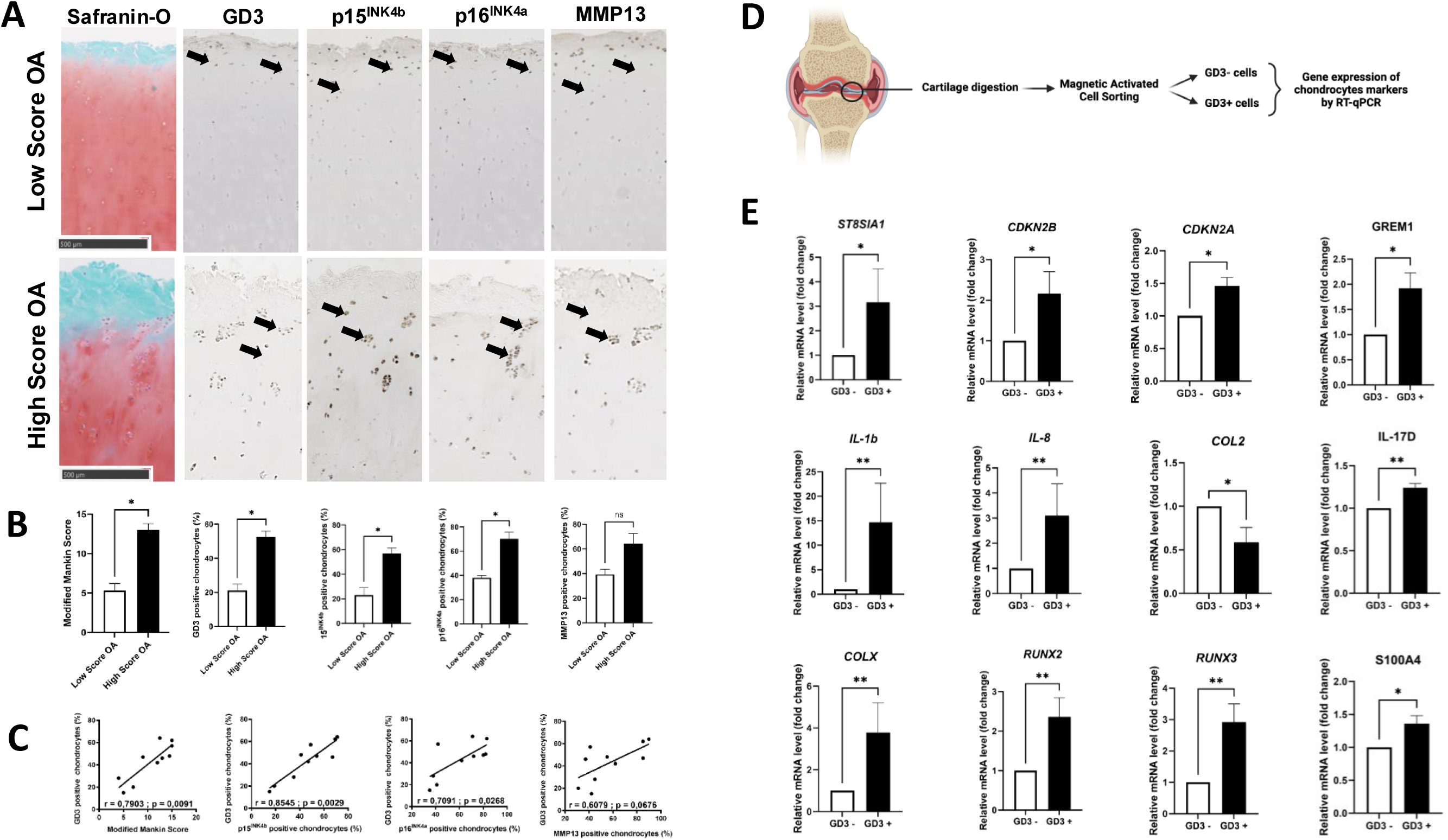
GD3 positive chondrocytes correlate with clinical score in human OA pathogenesis and express senescence markers. **A)** Experimental design of isolated fresh chondrocytes for flow MACS. **B)** Gene expression analysis of senescence and hypertrophic markers by RT-qPCR in GD3 enriched chondrocytes compared to depleted chondrocytes (n=7). * p < 0,05. Statistical significance was determined by unpaired, two-tailed Student’s t-test. **C)** Representative images for the tibial plateau of Safranin-O-fast green staining of low score OA (n=3) and high score OA (n=7) samples. The quantification of degradation OA Score was performed using Mankin Score. **D)** Representation immunohistochemical (IHC) staining for p15^INK4b^, p16INKAa, p21, MMP13 and GD3, with quantification of positive cells performed with QuPath-0.3.2. (n=10) **E)** Correlation between GD3 positive cells, OA Score and senescence hallmarks positive cells. The scale bar indicates 500 µm. *p < 0,05. Statistical significance was determined by unpaired, two-tailed Student’s t-test.

In addition, to confirm the link between GD3 expression and senescence onset in other joint compartment, we also sorted GD3-positive and GD3-negative synoviocytes from freshly collagenase type IV digested OA patient synovium (Supplementary ***Fig. S2a***). By cytometry, the ganglioside GD3 was confirmed on the surface of OA synoviocytes (Supplementary ***Fig. S2b***). RT-qPCR analysis of GD3-positive and GD3-negative synoviocytes showed again higher expression of *ST8SIA1* and senescence -associated markers (*CDKN2B*, *CDKN2A*, *IL6* and *MMP3)* in GD3-positive than in GD3-negative cells (Supplementary ***Fig. S2c***). In conclusion, two different GD3-positive cell-types from the OA joint are enriched in senescence features.

### GD3 accumulation correlates also with the presence of p15^INK4b^- and p21-positive senescent cells and cartilage degeneration in an experimental murine OA-induced model

We then assessed *St8sia1* expression and GD3 accumulation in an animal preclinical model in which OA was induced by intra-articular injection of collagenase VII (CiOA) or 0.09% NaCl as sham control **(*Fig. 3a*)**. Following different days post-injection, mice were sacrificed and treated joints were harvested for either gene expression, subchondral bone morphometric analysis or histological scoring **(*Fig. 3a*)**. Gene expression analysis at day 42 showed the significant increase of *St8sia1* and senescence hallmark genes *Cdkn2b, Cdkn1a, Mmp13, Adamts5* expression in whole CiOA injected joints compared with the control sham joints injected with 0.9% NaCl **(*Fig. 3b*)**. Following collagenase injection in each treated joint, by microCT analysis, we also observed a subchondral bone remodeling as revealed by an increase in bone surface on bone volume ratio (BS/BV) compared to sham joint ***(Fig.3c)***. Furthermore, using Safranin-O/Fast green staining of harvested joints, we followed and classified cartilage degeneration based on the modified OARSI score [14]. We could distinct 3 groups (<5 for healthy, 10-20 for medium and 20-30 high score) during the time course of the experiment with higher degrading score at day 28 and day 42 compared to sham controls (***Fig. 3d*)** [14]. In order to access articular GD3 accumulation and senescence markers expression in function of progressive joint remodeling, we determined by IHC the expression level of GD3, and p15^INK4b^, p16^INK4a^ or p21 and the SASP factor MMP13 on paraffin-embedded articular samples at day 14, day 28 or day 42 **(*Fig. 3d*)**. We found a significant correlation between cartilage degeneration and GD3-positive chondrocytes (r=0.6667; p=0.0209), as well as with p15^INK4b^-(r=0.8491; p=0.0008) or p21-positive chondrocytes (r=0.833; p=0.0012) (***Fig. 3e*)**. In contrast, a weak non-significant correlation was found with p16^INK4a^ and MMP13 positive chondrocytes confirming previous findings by others [26]. Remarkably, as found in human samples, the percentage of GD3-positive chondrocytes correlates also with the percentage of p15^INK4b^ (r=0.721; p=0.0116), as well as with p21-positive cells (r=0.933; p<0.0001) or MMP13 (r=0.621; p=0.0319) but not with p16^INK4a^ positive chondrocytes **(*Fig 3f*)**. In addition, analysis by immunostaining of the synovial membranes on the same murine joint samples showed that GD3-positive cells are co-detectable with senescence markers in high synovitis score (>2) **(*Supplementary Fig. S3*)**. These *in vivo* findings confirmed that, in mice as in patients, cellular senescence progressively increases during OA severity, and that the number of GD3-positive cells increases in parallel with the joint degradation and with the expression of known senescence markers.

**Figure 3:**
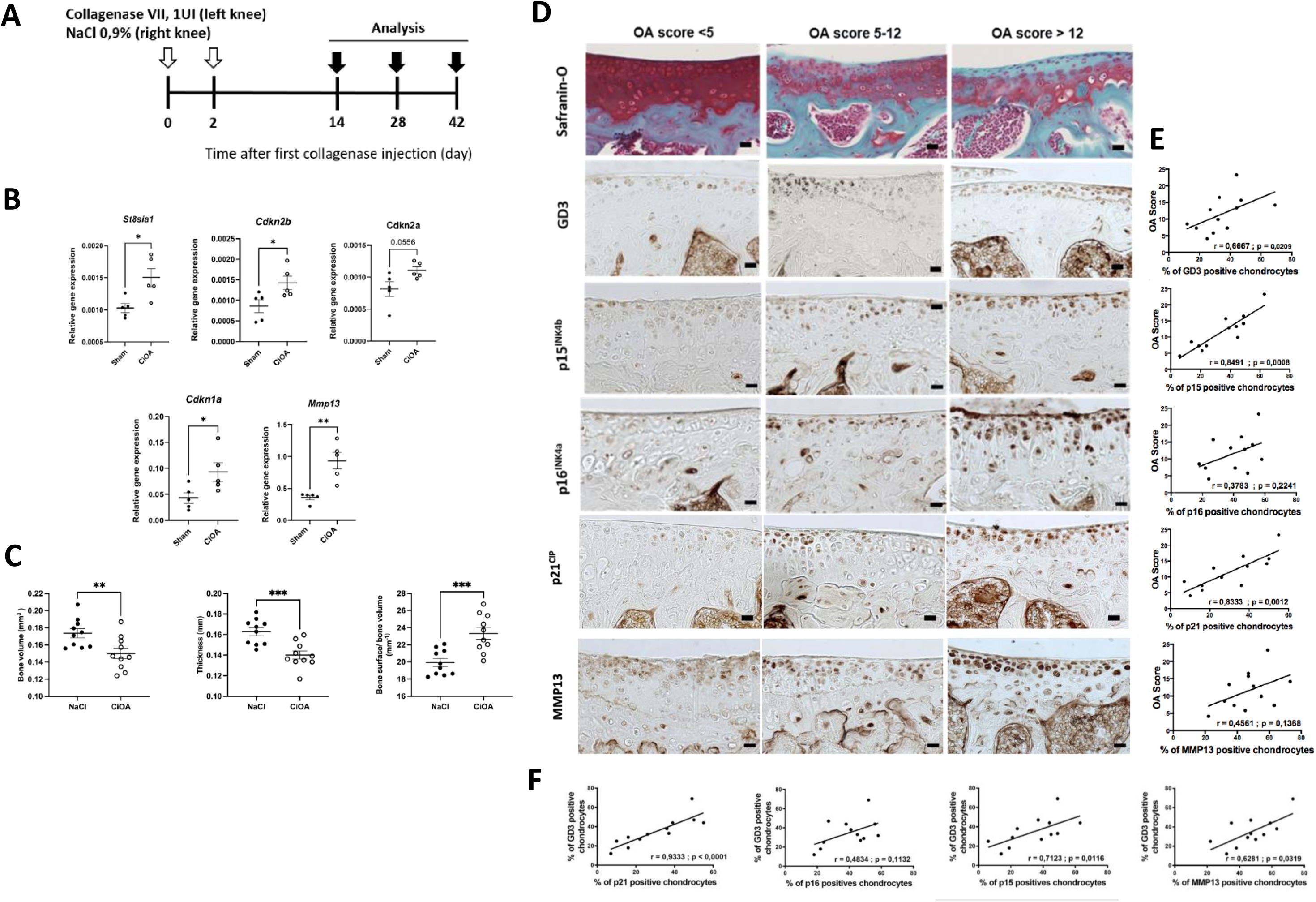
*St8sia1* gene and its product GD3 are induced in experimental murine OA model. **A)** Experimental design of the collagenase-induced osteoarthritis (CiOA) at day 0 and 2 with intra-articular collagenase VII (5U) or sham injection in 3-month-old C57BL/6 male mice. Gene and bone analysis were done at day 42. Cartilage analysis was performed at day 14, 28 or 42 after sacrifice. **B)** Gene expression of whole treated joint (control sham vs CiOA) was analyzed by RT-qPCR, n=5 samples per group. **C)** Analysis of the bone volume, bone thickness and bone surface/bone volume (BS/BV) by microCT between sham and CiOA knee (n=10). **D)** Safranin-O/Fast Green and Immunohistochemical staining of GD3, p15^INK4b^, p16^INK4a^ and MMP13, represented in the OA cartilage. Bar = 250 um. **E)** Graphical representation of Spearman correlation (r) test that was performed between OA Score, GD3 and senescence associated markers. **F)** Spearman correlation between GD3 positive chondrocytes and chondrocytes expressing senescence markers. All experiments are performed with n=12 mice. P-values are calculated by the Mann-Whitney test (unpaired t test, two-tailed). P-values are calculated by the Mann-Whitney test (unpaired t test, two-tailed).

### Intra-articular injection of anti-GD3 monoclonal antibodies increases sub-chondral bone volume and thickness in the experimental murine OA model

We have recently published that intra-peritoneally administered anti-GD3 monoclonal antibodies have beneficial effects on several age-related tissue phenotypes and associated senescence in naturally aged 21-month-old wild-type mice [11]. We therefore tested the therapeutic potentials of an anti-GD3 antibody in experimentally-induced murine OA model through local articular delivery. For this, we relied on the same CiOA model in which articular senescence and GD3 were detected. We injected a blocking monoclonal antibody against GD3 (or isotype IgG3 control) into the CiOA joint twice, at day 7 and 11 after OA induction before the peak of detectable senescence [27] (***Fig. 4a***). After sacrificed at day 42, RT-qPCR analysis of whole joints showed that the expression of *St8sia1*, *Cdkn1a, Cdkn2b, Mmp13, Adamts5* and *Tnfα* were decreased in the group treated with the anti-GD3 antibody compared with the isotype control group (***Fig. 4b***). Thus, we confirmed that this anti-GD3 monoclonal antibody can target cells expressing senescence-associated markers. In addition, we analyzed the cartilage, synovial membrane and subchondral bone structures of treated joints by confocal laser scanning microscope (CLSM), immunohistochemistry and microcomputed tomography, respectively. CSLM did not highlight any difference in cartilage 3D structure between groups (***Fig. 4c***). Similarly, the activation of the synovial membrane did not decrease, as shown by the synovitis score (***Fig. 4d***). Conversely, the anti-GD3 antibody induced a significant increase in bone volume and thickness in treated joints as well as a decrease in the bone surface/bone volume ratio confirming its positive effects on the subchondral bone structure compared to IgG control injection (***Fig. 4e***). Importantly, injection of the anti-GD3 or isotype control antibody in the sham non-OA joint (same doses and days) did not have any effect on bone remodeling, cartilage and synovial, indicating that this effect of the anti-GD3 antibody is only linked to the collagenase-induced pathological environment in CiOA joint (***Supplementary Fig.S4***).

**Figure 4:**
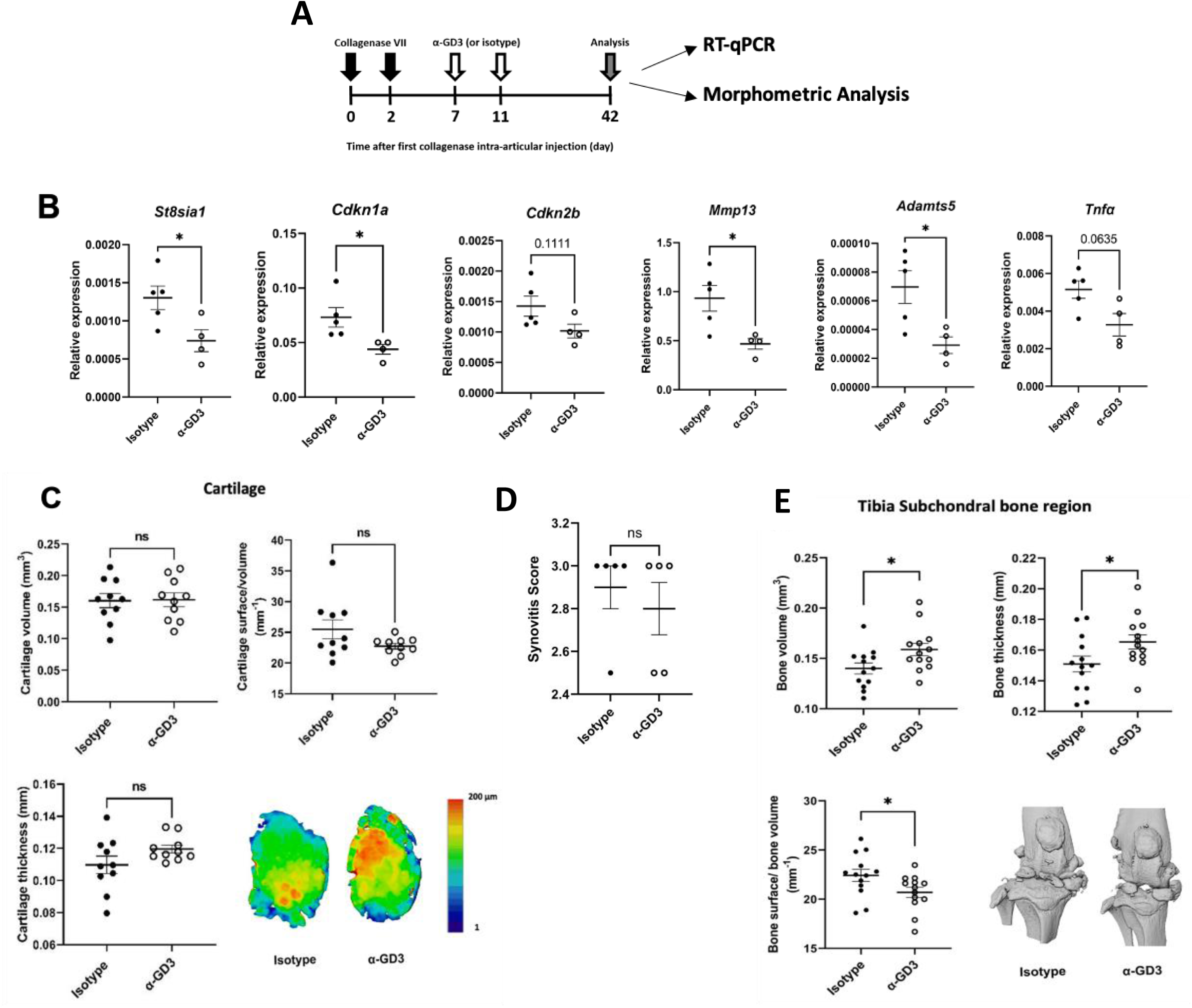
Effect of intra-articular anti-GD3 monoclonal antibody injection in an experimental murine OA model. **A)** Experimental design of the collagenase-induced osteoarthritis (CiOA) at day 0 and 2 with intra-articular collagenase VII (5U) injection in 3-month-old C57BL/6 male mice. Analysis was performed at day 42 after sacrifice. **B)** Gene expression analysis from whole treated joint was analyzed by RT-qPCR, n=4/5 simples per group. **C)** Confocal laser scanning microscopy, 3D reconstruction of cartilage. Analysis of the cartilage thickness, volume and ratio surface/volume (S/V) between isotype knee (n=10). **D)** Synovial Score by IHC (n=5/per group) **E)** High-resolution imaging of bone structure (microCT). Analysis of the bone volume, bone thickness and bone surface/bone volume (BS/BV) between Isotype and α-GD3 knee (n=15). P-values are calculated by the Mann-Whitney test (unpaired t test, two-tailed).

### Bone remodeling in OA pathogenesis may involve increased expression of GD3 synthase, senescence and functional markers in subchondral osteoclasts

On the previous figure, we revealed that an immunotherapeutic strategy targeting GD3 modulates bone structure in murine OA model. Bone homeostasis is the result on the balance between bone formation by osteoblasts and the catabolic functions of osteoclasts [28]. During aging or OA progression, bone and in particular subchondral bone homeostasis, is compromised, mainly due to an increase in osteoclast numbers and activity, in parallel with reduced bone formation so-called osteoblastogenesis [28]. Moreover, GD3 and its precursor GM3 were known to play a role in bone homeostasis, as demonstrated in mice in which knockout of *St8sia1* gene results in significant protection against age-related bone loss [9]. As this study proposed that GD3 effects on bone homeostasis could be explained by a change in the proportion of osteoclasts, we determined the proportion of osteoclasts (TRAP-positive cells) in the subchondral bone of anti-GD3 treated group compared to isotype treated group. We did not observe any significant difference in the number of TRAP-positive cells in both groups neither in gene expression of osteoblast makers (data not shown) (***Fig.5a***). Therefore, to explain such observed increase in bone volume, the anti-GD3 antibody may instead modulate the bone resorption activities of osteoclasts. To support such hypothesis, we monitored by RT-qPCR changes in functional osteoclastic gene expression in treated joints. We found that 3 canonical markers associated with osteoclast-dependent bone remodeling activities namely *Acp5, Mmp9* and *Ctsk* which encode respectively for TRAP degrading activity, for MMP9, an organic bone matrix degrading enzyme, and for Cathepsin K, a highly specific osteoclast catabolic enzyme, were all significantly downregulated in the anti-GD3 treated group compared with the isotype group (***Fig. 5b***).Altogether these findings argue for a role of GD3 in controlling genes-related to osteoclast activities in OA conditions as found previously during aging [9]. To confirm a specific increase expression of GD3 synthase and senescence markers in osteoclast during OA progression, we compared in kinetic (day 4 and 10) the *in vitro* osteoclastogenic differentiation profile of monocytes derived from bone marrow flush of mice injected with collagenase (CiOA) versus injected with NaCl. We observed that, in response to M-CSF+RANK-L cocktail, differentiated osteoclasts from CiOA mice have a significant increased expression in *St8sia1* enzyme as well as the senescence marker *Cdkn2a* compared to NaCl at day 4 and acquired their functional markers more rapidly confirming the increase osteoclastic activities in OA murine model (***Fig. 5e***). Finally, to confirm such findings during human OA pathology, we investigated whether *ST8SIA1 and CDKN1A or CDKN2A* gene expression was also increased in subchondral bone marrow isolated from OA patients compared to healthy donors. For that, we applied again the previous k-mer method on dedicated publicly available RNA seq data [29] (***Fig. 5c***). We observed that *ST8SIA1* expression was increased altogether with an increase in *CDKN1A* and markers associated with osteoclastic activities such as *ACP5, CTSK* and *MMP9* in the OA group compared with the healthy control group (***Fig. 5d***). These results confirmed that OA-induced pro-senescent micro-environment triggers GD3 synthase expression and favors osteoclast activities in mice as well in OA patients. Targeting GD3 can reduce such deleterious effects on bone homeostasis.

**Figure 5:**
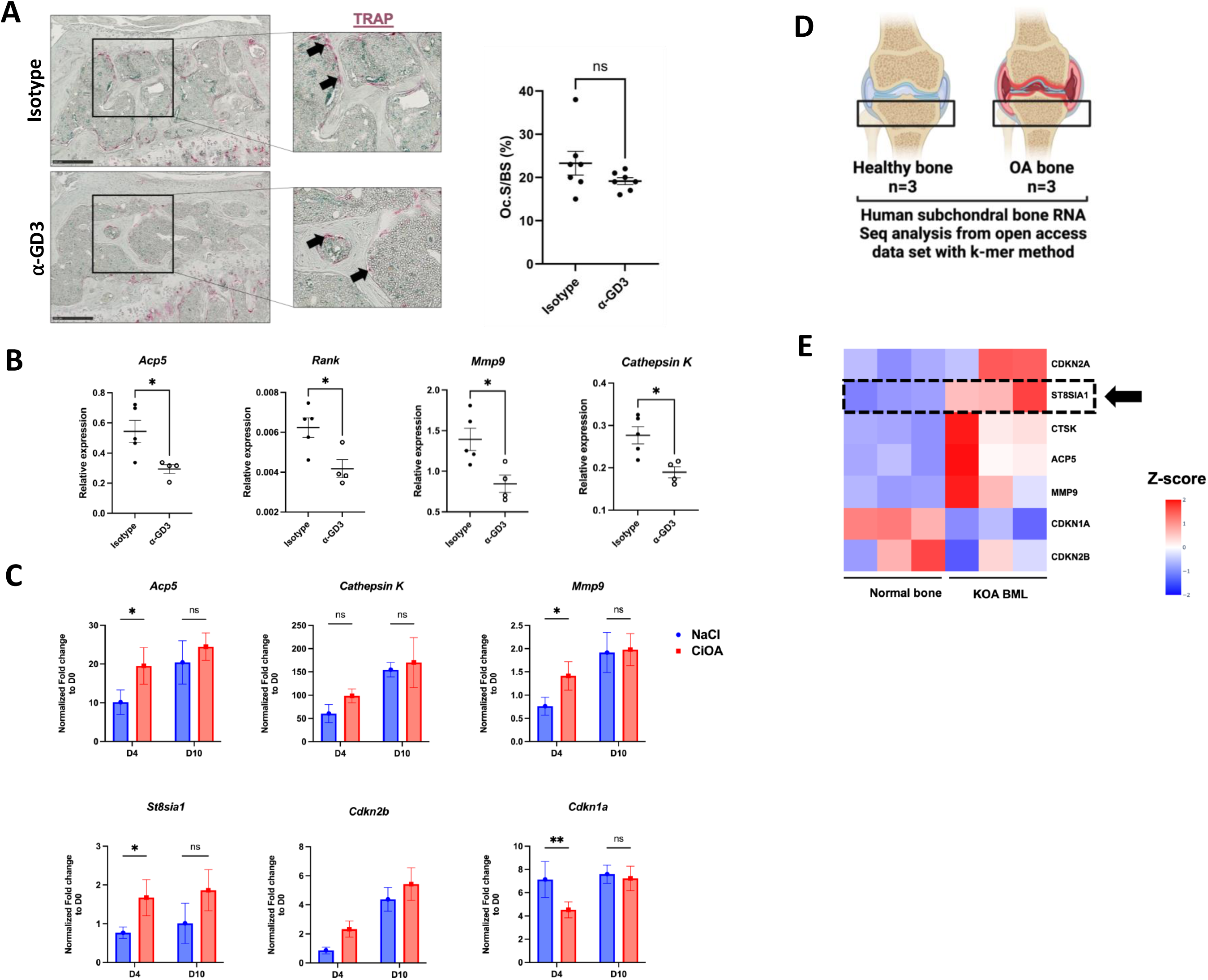
GD3-positive osteoclasts may be responsible for bone remodeling in the OA condition. **A)** Representative images of tartrate-resistant acid phosphatase (TRAP) staining and quantification of TRAP positive multinuclear cells. The scale bar indicates 250 µm. n=7. **B)** Osteoclasts gene expression analysis from whole treated joint was analyzed by RT-qPCR, n=4/5 simples per group. *p < 0,05, Statistical significance was determined by unpaired, two-tailed Student’s t-test. **C)** Isolated bone marrow monocytes/macrophages were treated with M-CSF and RANKL to obtain mononuclear preosteoclasts (day 4, D4) and mature osteoclasts (day 10, D10), from CiOA or NaCl injected mice at day 42. Normalized data compared to day 0. * p < 0,05, Statistical significance was determined by Two-way ANOVA (n=3). **D)** Experimental design from RNA-Seq analysis, bone/subchondral bone from normal versus OA Bone marrow lesion. **E)** Heatmap represented transcriptomic signature with hallmarks of pathology in bone.

## DISCUSSION

In this study, we investigated the correlation between expression of ganglioside GD3 or its synthase (encoded by the *ST8SIA1* gene) with OA-related senescence. GD3 expression at the cell surface of OA chondrocytes was associated with cellular senescence as confirmed by an increase in *CDKN2B* and *CDKN2A* but also SASP (*IL6*, *IL8*, *IL1β, ADAMTS5* and *MMP13*) markers in sorted GD3-positives cells. Remarkably, GD3 synthase and GD3 expression was also increased in a 3D organotypic OA senescence cartilage model confirming this direct link. Remarkably, using immunostaining approaches, we have shown that both GD3 and the senescence markers are associated with severity of cartilage degeneration in OA patients but also in a preclinical OA mouse model. It would be interesting in the future to determine whether this association between GD3 and loss in cartilage structure is maintained with the common clinical score, meaning the Kellgren Lawrence (KL) classification, which is based on radiographic severity data of OA patients [30]. Limitation of such studies is always that the cartilage samples come from prosthetic surgery on patients with high KL score.

Among the various others tissue found in the OA joint, there is the activated inflamed synovium. In OA-derived synovium, we could reveal by scRNA seq analysis that *ST8SIA1* is exclusively expressed by synovial-derived fibroblasts, specifically two distinct subgroups, S2 and S3 that can be differentiated by the expression of *CXCL12* (a pro-inflammatory marker) [31] and periostin, *POSTN* (associated with pathogenic pathways) respectively [13]. As found in cartilage, sorted GD3-positive synoviocytes expressed also high level of senescent markers compared to GD3-negative synoviocytes. It would be interesting to assess whether GD3 expression follows the increase in the percentage of p16^INK4a^-expressing synoviocytes that accumulate in inflamed OA synovium [32,33]. Overall, our results show that senescent OA chondrocytes and senescent OA fibroblast-like synoviocytes express the GD3 synthase and GD3 at the cell surface. This makes of GD3 level a strong indicator of senescence in OA joints.

Senescent cells, which accumulate in joint tissues throughout lifespan, are involved in the development and progression of OA. Many research groups have explored the therapeutic potential of targeting senescent cells by selecting drugs for clinical trials in OA patients [34]. In this study, we investigated the senotherapeutic effects of a neutralizing anti-GD3 antibody injected into the joint of a preclinical mouse model of OA. We showed that targeting GD3 had a significant protective effect only in the subchondral bone compartment, but not on cartilage or on synovitis score. This relatively disappointed result could be explained by several factors. The short lifetime of the antibody in the joint but also the limited accessibility to articular senescent cells due to joint composition in proteo-aminoglycans, are the main limitations [35]. The development of dedicated nanomedicines for articular delivery is the next challenge of the field. Nevertheless, this partial senotherapeutic effect observed is reminiscent of other published data [34]. These results could be due to the heterogeneity in senescent cells and the lack of clear characteristics for each subpopulation in OA joints.

Importantly, this study confirm our recent published observation for the positive protective effects of the anti-GD3 antibody on the spontaneous subchondral bone remodeling observed in aged mice [11]. Indeed, during the course of OA disease or aging process, osteoclasts play a critical role in joint remodeling by increasing bone resorption [28]. The excessive number and activity of osteoclasts disturbs the subchondral bone balance and contributes to joint destruction [36]. We observed a significant reduction in the expression of genes that regulate osteoclast resorption activities (*Acp5, mmp9* and *Csk*) following anti-GD3 antibody administration in the CiOA joint (Fig 6). This finding suggests that targeting GD3 leads to a reduction in osteoclast activity in CiOA mice and importantly, not in sham mice. It is worth noting that the GD3 knock-out mouse has a reduced number of osteoclasts, suggesting a role for this ganglioside in osteoclastogenesis. Remarkably, GD3 is also known to enhance adhesion signals to extracellular matrix by regulating integrins binding properties of melanoma cells [37]. One can then speculate that OA-induced enrichment in GD3 at the cell surface of osteoclasts near to integrins that are essential for their bone-binding activities will favor osteoclast attachment and increase their lytic activities against bone structure. Thus, we propose that the injected neutralizing antibody could interfere with such processes reducing the ability of osteoclasts to anchor and erode bone, thus providing protection against bone remodeling in OA conditions. Further experiments combining *in vitro* osteoclastogenic bone erosion assays in presence of an anti-GD3, are needed to confirm these claims. Finally, we found that human *ST8SIA1* and *CDKN1A* genes were also increased in subchondral bone in OA patients compared with the control group confirming that GD3-positive cells are accumulating with senescence and osteoclastogenic markers during disease progression in mice and human.

A recent study showed that CD26, a surface marker encoded by the dipeptidyl peptidase (*DPP4*) gene, is also a hallmark of senescence in OA cartilage [30]. CD26-positive OA chondrocytes shared common features with GD3-positive OA chondrocytes, such as upregulation of some senescence or catabolic markers and downregulation of a chondrogenic marker (*COL2*). In the future, we could investigate the correlation between GD3 and CD26 expression in view of developing dedicated bispecific therapeutic tools to target senescent joint cells. In addition, as the number of GD3-positive and CD26-positive cells in OA joints increases with tissue degradation [30], short-lived radiolabeled bi-specific anti-GD3/CD26 antibodies could become valuable tools for assessing the level of joint senescence in OA patients by non-invasive bio-imaging. Indeed, with PET imaging after articular injection, these tools could be used for OA prognosis and appropriate patient stratification based on their level of joint senescence. This bi-specific immune-tool could even help to assess the clinical efficacy and efficiency of future senotherapeutics as a companion test to monitor recovery of OA patient in clinical trials.

To conclude, the results of our study propose that the ganglioside GD3 as novel cell surface marker of OA-related senescent phenotype in various joint compartments under osteoarthritic conditions. The development of dedicated immune-tools targeting GD3 could be useful for clustering OA patients based on their level of senescence but also for delaying OA-induced subchondral bone remodeling as innovative therapeutic approaches in joint treatment.

## Acknowledgments

We deeply thank Léa Marinèche for technical supports on osteoclast staining, Florence Apparailly for critical reading of the manuscript and the Montpellier University Hospital services for sharing human samples. We thank E-Cell France and the RHEM platform for bone morphometric and histological analysis, Anne-Laure Bonnefont’s supports and the animal facilities RAM platform. CJ, EG, JC, YMP and JMB received funds from INSERM National Ageing Network (AGEMED), Montpellier University and the French association on calcified tissue, OSCAR. CF was recipient of a PhD studentship from FHU/CHU RegenHab and Montpellier University. GM is a PhD fellow from INSERM and Montpellier University Hospital. Biorender license was owned by CF.

## Authors’ contributions

CF, GM, YMP and JMB conceived the study, designed, and supervised the experiments. CF, MM, GM, BG, RS, TC, LD, performed the experiment and analyzed data. EG, JC provided the therapeutic anti-GD3 antibody and the concept of targeting GD3. LD, RFL and YMP provided signed patient consents and joint explants from OA patients. CF, GM, YMP and JMB drafted the article. CJ, EG, JC, YMP and JMB provided financial support.

## Abbreviations

GD3: ganglioside 3
SASP: secretory associated senescence phenotype
CKI: Cyclin-dependent kinase inhibitors
CiOA: Collagenase induced Osteoarthritis
POSTN: Periostin
IHC: immunohistochemistry
microCT: micro-Computed Tomography

---

**Figure S1:**
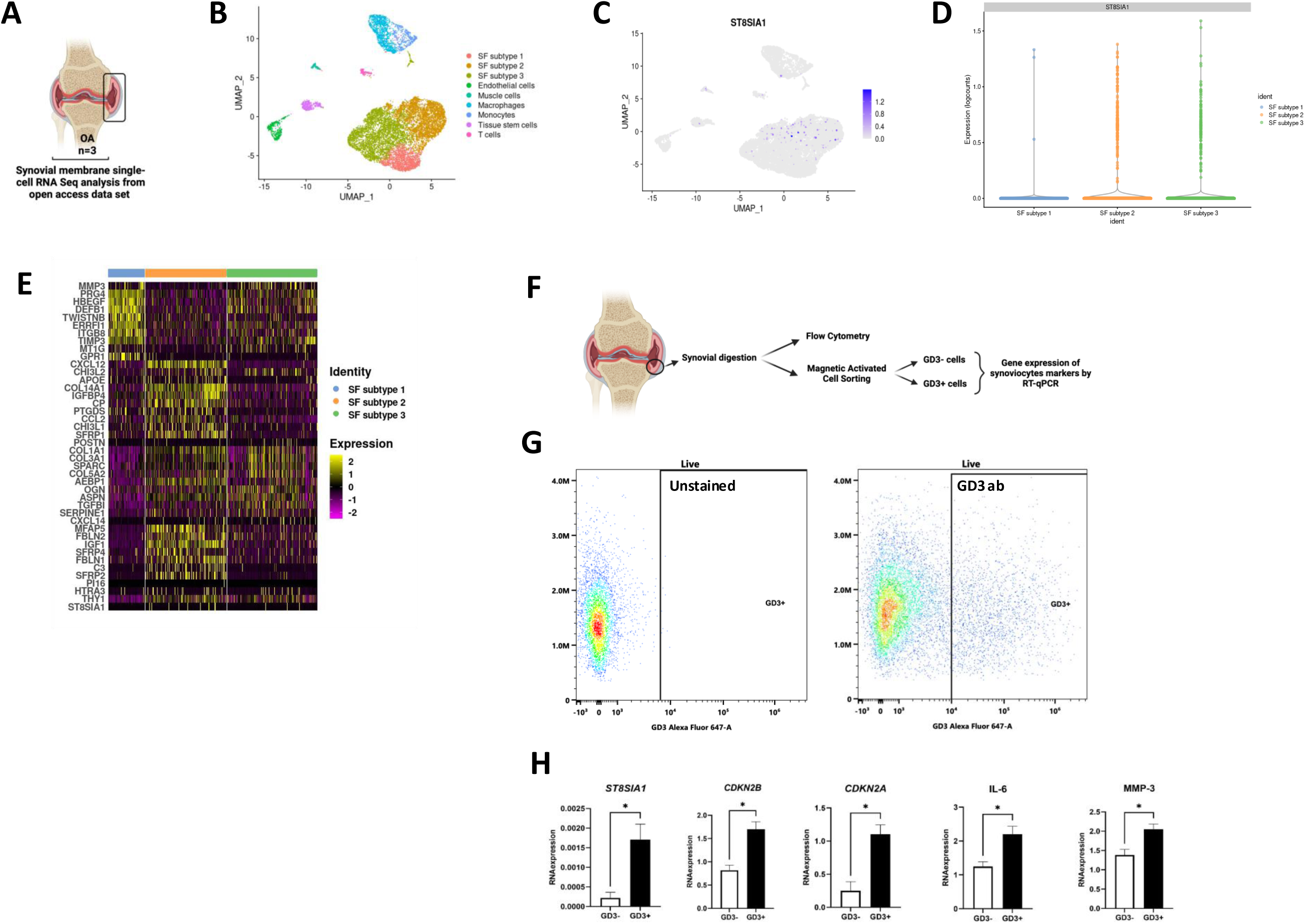
Human GD3positive fibroblasts expresses senescence markers in OA pathogenesis. **A)** Experimental design from RNA-Seq single cell analysis. **B)** Uniform manifold approximation and projection (UMAP) plot of scRNA-seq show unsupervised clusters colored according to putative cell types among synovial-fibroblast (SF) subtype 1, synovial-fibroblast (SF) subtype 2, synovial-fibroblast (SF) subtype 3, Endothelial cells, Muscle cells, Macrophages, Monocytes, Tissue stem cells and T cells. **C)** *ST8SIA*1 expression in UMAP plots. **D)** Violin Plot and **E)** Heatmap of unsupervised clustering analysis shows the top ten highly expressed genes per synovial-fibroblast (SF) subtypes as determined by Seurat analysis. Expression level is scaled based on z-score distribution. **F)** Experimental design of isolated fresh fibroblasts for flow cytometry and MACS. **G)** Representative flow cytometry showing GD3 positive cells in osteoarthritic synovium **H)** Comparative gene expression analysis of senescence and hypertrophic markers by RT-qPCR in GD3 enriched synoviocytes compared to GD3-depleted ones (n=7). * p < 0,05. Statistical significance was determined by unpaired, two-tailed Student’s t-test.

**Figure S2:**
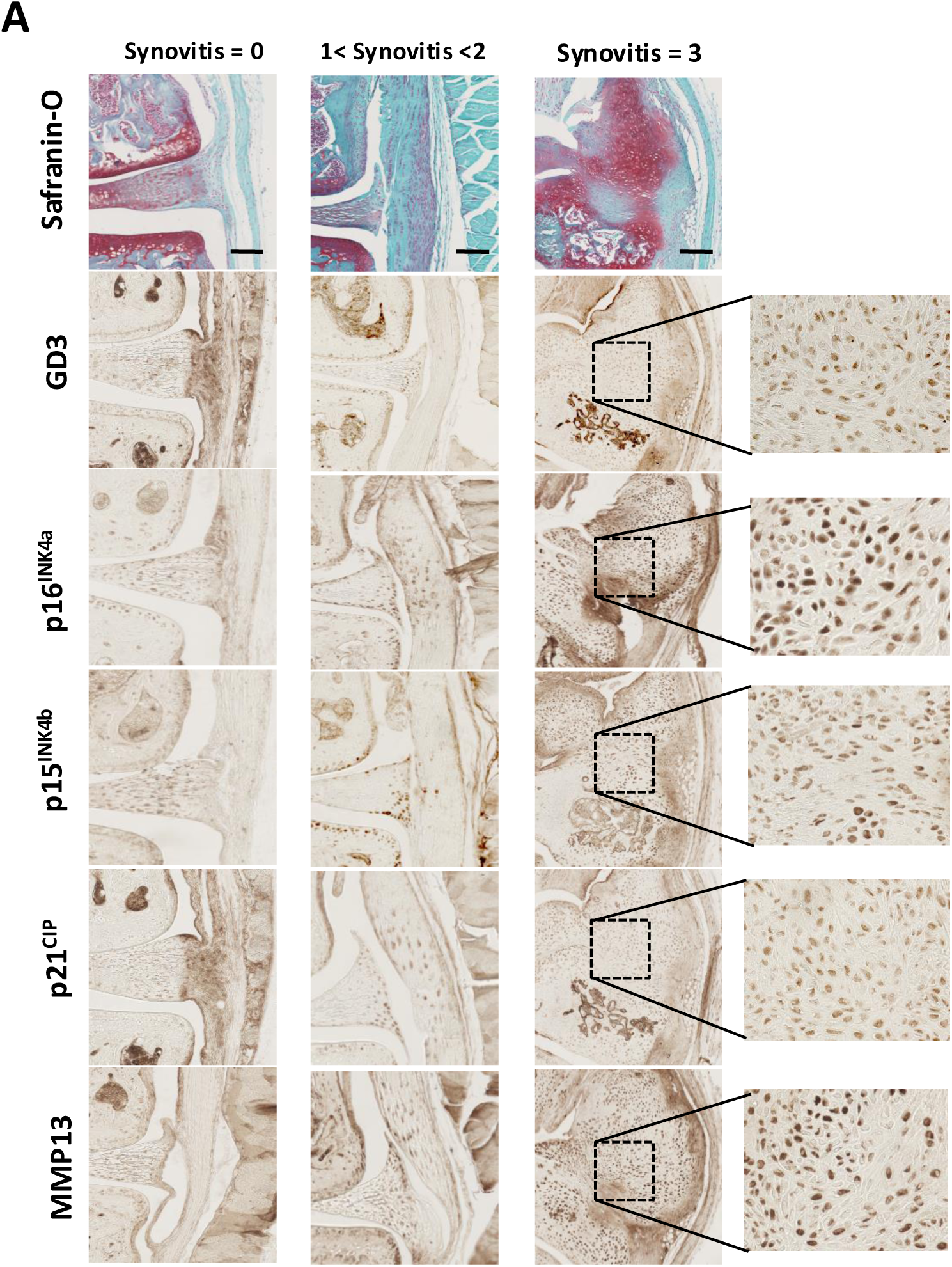
Presence of GD3 and markers associated to senescence in CiOA murine synovium. Safranin-O/Fast Green and Immunohistochemical staining of GD3, p16^INK4a^, p15^INK4b^, p21^CIP^ and MMP13, represented in the OA synovium. Bar = 250 um. All experiments are performed with n=12 mice.

**Figure S3:**
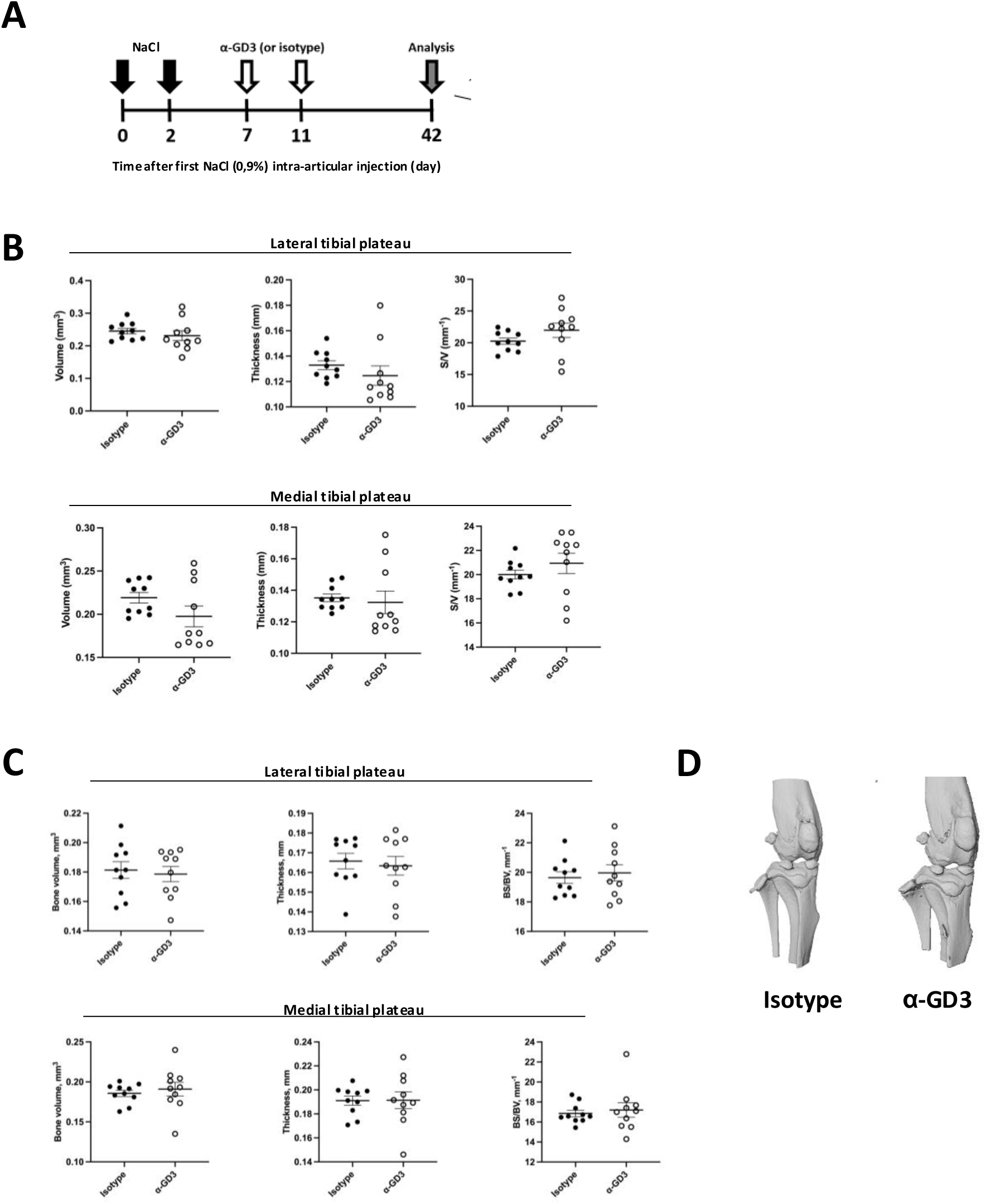
Effect of intra-articular anti-GD3 monoclonal antibody injection in an experimental murine OA model. **A)** Experimental design of the sham group at day 0 and 2 with intra-articular physiological serum (NaCl 0.9%) injection in 3-month-old C57BL/6 male mice. Analysis was performed at day 42 after sacrifice (n=15 mice per group). **B)** Confocal laser scanning microscopy, 3D reconstruction of cartilage. Analysis of the cartilage thickness, volume and ratio surface/volume (S/V) between isotype knee. **C)** High-resolution imaging of bone structure (microCT). Analysis of the bone volume, ratio bone surface/bone volume (BS/BV), bone thickness and ratio bone surface/tissue volume (BS/TV) between isotype knee and α-GD3 knee. Statistical significance was determined by unpaired, two-tailed Student’s t-test.

